# A novel genomic DNA library preparation method with low GC bias

**DOI:** 10.1101/2022.01.28.478268

**Authors:** S. Thomas Kelly, Tsuneo Hakoyama, Kie Kumaishi, Haruka Okuda-Yabukami, Sachi Kato, Makoto Hayashi, Aki Minoda, Yasunori Ichihashi

## Abstract

The amount of input DNA available to prepare next-generation sequencing (NGS) libraries is often limited, which can lead to GC content bias and enrichment of specific genomic regions with currently available protocols. In this study, we used breath capture technology to incorporate sequencing adapters into DNA to develop a novel cost-effective protocol for the preparation of genomic DNA libraries. We performed a benchmarking experiment comparing our protocol with common commercially available kits for genomic DNA library preparation with input DNA amount in the range of 1 to 50 ng. Our protocol can generate high-quality genomic sequence data with a marked improvement in coverage breadth and low GC bias, in contrast to standard protocols. Further, our protocol reduces sample handling time and reagent costs, and requires comparatively fewer enzymatic steps relative to other protocols, making it suitable for a range of genomics applications.

## Introduction

Next-generation sequencing (NGS) platforms have been used for a wide range of investigations in genomics and related applications. However, several library preparation methods require a high amount of input DNA, which may not be feasible for some applications (Li *et al.*, 2019). For example, very low DNA yields are common in analyses such as chromatin immunoprecipitation (ChIP-seq; commonly performed for epigenomic analyses), degraded DNA from field samples (for example in forensic science, archaeology, and paleontology) (Gansauge and Meyer, 2013; Xavier and Parson, 2017), parasites and microorganisms that cannot be cultured (Nascimento *et al*., 2016), viruses (Malboeuf *et al*., 2013), very small insects (Cruaud *et al.*, 2019), and metagenomics of environmental samples (Rinke *et al.*, 2014). Low DNA input can also be an issue in single-cell analyses (Harada *et al.*, 2019; Kaya-Okur *et al.*, 2019), analyses using circulating fetal or tumor DNA (Mauger *et al.*, 2020), and clinical studies with formalin-fixed paraffin-embedded tumor tissue samples (Munchel *et al*., 2015; Kader *et al.*, 2016; Zhang *et al.*, 2019).

Commercially available kits, such as TruSeq® Nano (Illumina), Nextera® (Illumina), and ThruPLEX® (Takara Bio) are popular for library preparation for NGS experiments with low sample input. TruSeq (Illumina) kits have been shown to produce high-quality data for RNA-Seq using low cDNA input compared to other protocols (Song *et al.*, 2018; Sarantopoulou *et al.*, 2019). TruSeq Nano remains one of the most popular and reliable library preparation techniques (Li *et al*., 2019; Pasquali *et al*., 2019) using a low sample input (as low as 1 ng) and higher sensitivity than TruSeq version 2 (Rhodes *et al*., 2014) or NEBNext® Ultra. The Nextera genomic DNA library preparation kit that uses the Tn5 transpose can be used to prepare libraries rapidly using a low input at 1 ng or lower with high sensitivity. However, the kit is costly and shows GC coverage and sequence biases compared to the TruSeq kits (Adey *et al*., 2010; Lan *et al*., 2015; Sato *et al*., 2019). Nextera kits are also susceptible to the presence of contamination or duplicates (Rinke *et al*., 2014), do not detect more variants than lower-cost alternatives (Pasquali *et al*., 2019), and the limited control of fragment size can introduce biases (Adey *et al*., 2010; Nascimento *et al*., 2016). The Rubicon Genomics ThruPLEX kit (Takara Bio) has been shown to have relatively high sensitivity for *de novo* genome assembly and reliably detects variants in low-input DNA (Chung *et al*., 2016; Nascimento *et al*., 2016; Mauger *et al.*, 2020) and ChIP-Seq samples (Sundaram *et al.*, 2016).

Because library preparation can be expensive, there have been several efforts to reduce the cost, including dilution of kits (Cruaud *et al.*, 2019; Li *et al.*, 2019) and utilization of cheaper alternatives (Dunham *et al.*, 2019; Pasquali *et al.*, 2019). Notably, it is possible to reduce the cost of preparation of genomic and transcriptomic libraries without compromising the variant and gene detection rates or data quality (Combs and Eisen, 2015; Pasquali *et al*., 2019). Numerous protocols have been developed to enable ChIP-Seq with a low input (Schmidl *et al.*, 2015; Kaya-Okur *et al.*, 2019; Handa *et al.*, 2020). For example, ChIP-Seq studies in plant tissues would not be possible without these low-input techniques (Birkenbihl *et al.*, 2017; Zheng and Gehring, 2019). As such, the community would benefit tremendously from a simple, fast, and low-cost alternative preparation technique.

The cost of library preparation can be reduced significantly using custom protocols compared to commercial kits (Kumar *et al.,* 2012; Baym *et al.*, 2015). Previously, we have demonstrated the utility of the Breath Adapter Directional sequencing (BrAD-seq) protocol for the preparation of RNA-Seq libraries (Townsley *et al.*, 2015; Ichihashi *et al.*, 2018). The BrAD-Seq technique is a streamlined, ultra-simple, and fast library preparation method that uses breath-capture technology to incorporate strand-specific adapters, enabling large-scale transcriptomic studies at a lower cost. In this study, we adapted the breath capture technology in the BrAD-Seq protocol (shown in Fig. 1) to prepare low sample input libraries for DNA sequencing and compared the results obtained with commercial kits, namely Illumina TruSeq and Rubicon ThruPLEX. The revised protocol has the advantages of lower cost in consumables (Table 1) and a shorter protocol. A cheaper genome sequencing alternative for low input and poor-quality samples would expand the range of feasible applications in genomics and epigenomics experiments and increase sample sizes (Lakens, 2021).

**Table 1.** Cost per sample for each technique.

| Library | BrAD-Seq | TruSeq Nano | TruSeq ChIP | ThruPLEX |
| --- | --- | --- | --- | --- |
| Vendor |  | Illumina | Illumina | Takara Bio |
| Price (€) Euro | 2.36 | 44.31 | 78.75 | 52.07 |
| Price (¥) JPY | 304 | 5,708 | 10,146 | 6,709 |
| Price (\$) USD | 2.80 | 52.60 | 93.48 | 61.81 |

Prices are shown for Japanese distributors with exchange rates at the time of submission to indicate the relative costs alone. Exact costs depend on the distributors in each country.

**Figure 1.**
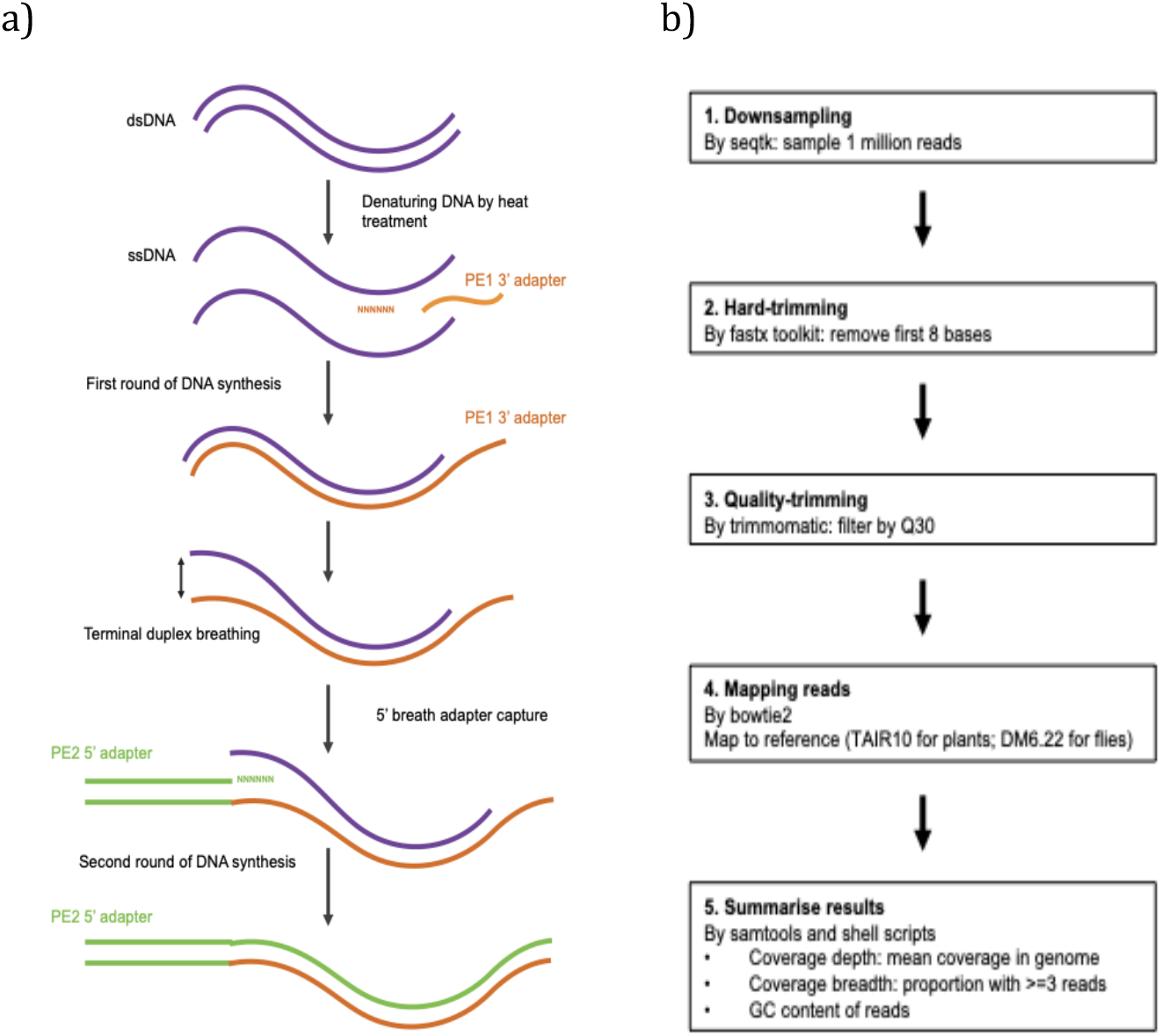
Experimental and bioinformatics workflows for BrAD-Seq using genomic DNA. a) Schematic representation of the breath capture technique for library preparation. Outline for how breath capture is performed to generate a library for next-generation sequencing from double-stranded DNA. b) Summary of steps in bioinformatics analysis. All samples were processed with the same parameters for comparison.

Protocols that reduce GC bias in libraries are necessary since variations in the GC content between genomes can cause biases in the analyses, requiring greater sequencing depth for *de novo* genome assembly (Kozarewa *et al.*, 2009; Benjamini and Speed, 2012; Chen *et al.*, 2013). GC bias can also be problematic for variant calling, metagenomics, and quantitative analyses by ChIP-Seq, as regions with low GC content will be under-represented in low-coverage libraries (Benjamini and Speed, 2012; Rinke *et al.*, 2016; Sato *et al.*, 2019; Browne *et al.*, 2020). Avoiding GC bias is also important when working with ancient DNAs, as it reduces the contamination of GC-rich microbial DNA (Dabney and Meyer, 2012). Amplification-free techniques have been developed previously to reduce GC bias; however, they require high amounts of input material, making GC bias a widespread issue in published data with low input amounts (Chen *et al.*, 2013).

Here we demonstrate the application of the BrAD-Seq method to genomic DNA by comparing it to commercially available kits by criteria such as coverage breadth and GC bias.

## Methods

### Sample materials and preparation

Genomic DNA samples were obtained from plants (*A. thaliana*) and fruit flies (*D. melanogaster*). Wild-type thale cress Col-0 genomic DNA (BioChain Institute Inc. BCH #D1634310-5) was purchased from Cosmo Bio (Tokyo, Japan). Genomic DNA was extracted from *D. melanogaster* (*w*^*1118*^ mutant strain BDGC #5905).

### DNA fragmentation

The plant genomic DNA samples were diluted with TE (10 mM Tris-HCl pH 8.0, 1 mM EDTA) to 10 ng/μl, and 20 μl of the diluted DNA samples were sheared by heat treatment (95°C, 45 min) into fragments ranging from 500-1000 bp. The fragmented DNA was purified by the addition of 30 μl Agencourt AMPure XP beads (Beckman Coulter) and mixed by pipetting. After 5 min, the samples were placed on a magnetic tray and the supernatants were removed. Pellets were washed twice with 80% ethanol. Residual ethanol was removed completely, and the pellets were air-dried for 2 min. The fragmented DNA samples were eluted with 10 μl of 10 mM Tris-HCl (pH 7.5) and quantified using a Nanodrop (Thermo Scientific). Samples were aliquoted into 1 ng, 10 ng, and 50 ng amounts. Samples were amplified by PCR with the cycling details described in Table S1 after preparation with each library protocol.

Fruit fly genomic DNA samples were prepared with the Focused-ultrasonicator S220 (Covaris; duty factor 10%, peak incident power(w) 140, cycle/burst 200, time 80 s) into fragments ranging from 300-400 bp. The samples were aliquoted into 1 ng and 10 ng aliquots and were amplified by PCR with the cycling details described in Table S2 after preparation with each library protocol.

### Generation of BrAD-Seq libraries for genomic DNA

The full BrAD-Seq protocol for genome DNA was deposited at protocols.io (https://dx.doi.org/10.17504/protocols.io.bsx9nfr6). The adapter and primer sequences used are listed in Table S3.

#### 3’ adapter priming

The fragmented DNA samples were aliquoted into different amounts (1 ng, 10 ng, and 50 ng for plants and 1 ng and 10 ng for flies) to evaluate the effect of DNA amount on the sequencing library quality. The DNA fragments were used as templates to produce libraries that had adapter sequences for Illumina sequencing at the 5’ and 3’ ends. After denaturing the double-stranded DNA (dsDNA) to single-stranded DNA (ssDNA) by heat treatment, the PE1 adapter sequence was added to the 3’ ends of the ssDNA using DNA polymerase in a 15 μl reaction mixture containing 1.5 μl 10 × Ex Taq buffer, 1.2 μl dNTPs (2.5 mM each), 0.075 μl (0.375 units) of Ex Taq (Takara), 5 μl of the fragmented DNA, 1 μl of 5 μM 3’ adapter oligo (L-3ILL-N8.2 in Table S3), and 6.225 μl H2O. The mixtures were incubated in a thermal cycler with the following program: 94°C for 2 min, 45°C for 10 min, 42°C for 10 min, 72°C for 5 min, and hold at 4°C. The synthesized DNA was purified by addition of 5 μl 50 mM EDTA (pH 8.0) and 30 μl Agencourt AMPure XP beads to each sample and mixed by pipetting. After 5 min, the sample was placed on a magnetic tray and the supernatant was removed. The pellets were washed twice with 80% ethanol. The residual ethanol was completely removed and air-dried for 2 min.

#### Breath capturing 5’ end of ssDNA

Introduction of the adapter sequence into the 5’ ends of ssDNA fragments was performed using breath capture technology with a 5’ double-stranded adapter oligo (Townsley *et al.*, 2015; Ichihashi *et al.*, 2018). The 5’ double-stranded adapter oligo was prepared by hybridization using 10 μl each of 10 mM oligos 5pSense8n and 5pAnti (described in Table S3) in 80 μl of H_2_O. The mixture was incubated in a thermal cycler with the following program: 94°C for 1 min; (94°C for 10 s) × 60 with −1°C/cycle, 20°C for 1 min, and hold at 4°C. The bead-bound DNA samples were eluted with 4 μl of 10 mM 5’ double-stranded adapter oligo. Subsequently, 6 μl of the following mixtures were added: 1 μl 10 × Pol I buffer, 0.25 μl dNTPs (25 mM each, Thermo Scientific), 0.25 μl DNA polymerase I (Thermo Scientific) and 4.5 μl H_2_O. The mixture was incubated at 25°C for 15 min. The samples were size-selected by the addition of 10 μl of 50 mM EDTA and 30 μl of ABR buffer (15% PEG 8000, 2.5 M NaCl) and mixed by pipetting. The samples were allowed to stand for 5 min and were subsequently placed on a magnetic tray. The supernatant was removed, and the pellets were washed twice with 80% ethanol. The pellets were air-dried for 2 min and resuspended in 30 μl of 10mM Tris-HCl (pH 7.5).

#### Enrichment and index sequence addition

Full-length adapter sequences were added to the libraries and amplified by PCR using the primers PE1 and PE2 (Table S3). The enrichment and adapter extension steps were performed in a reaction mixture containing 2 μl 10 × Ex Taq buffer, 1.6 μl dNTPs (2.5 mM each), 0.1 μl Ex Taq (Takara, cat# RR001A), 4.3 μl H_2_O, 10 μl breath captured DNA, 1 μl of 2 μM PE1 primer, and 1 μl of 2 μM PE2 primer. The reaction mixtures were incubated in a thermal cycler with the following program: 94°C for 2 min; 94°C for 30 s; 65°C for 30 s; 72°C for 30 s) × 16-21 (see Tables S1 and S2); 72°C for 7 min; and hold at 4 °C. The PCR products were cleaned with 16 μl Agencourt AMPure XP beads and the samples were mixed by pipetting. After 5 min, the samples were placed on a magnetic tray and the supernatants were removed. The pellets were washed twice with 80% ethanol. Residual ethanol was removed completely using a 10 μl pipette tip, and the sample was air-dried for 2 min. The libraries were eluted using 10 μl 10mM Tris-HCl (pH7.5).

### Generation of Illumina libraries using commercially available library preparation kits

TruSeq® Nano DNA LT Library Preparation Kit (Illumina) for plant samples and TruSeq® ChIP Sample Preparation Kit (Illumina) for fly samples were used according to the manufacturer's protocols. PCR cycling was performed as described in Tables S1 and S2. For the Rubicon Genomics ThruPLEX® DNA-seq Kit: High Performance Library Preparation for Illumina® NGS Platforms (currently available from Takara Bio USA), the manufacturer’s protocol was followed.

### Sequencing on the Illumina platform

The plant samples were sequenced on the Illumina MiSeq (single-end sequencing with 68 cycles and 8 bp index) with at least 1 million reads for each sample obtained.

The fruit fly samples were sequenced on an Illumina NovaSeq 6000 (paired-ends with 151 cycles and 8 bp index). Due to sample dilution factors, some samples were sequenced to a significantly greater depth than others.

### Bioinformatics analysis

A reproducible bioinformatics pipeline was developed in-house to ensure that all samples were processed using the same parameters. These were all run on the same architecture using a Debian 9 Linux server with an SGE job scheduler.

Samples were demultiplexed using bcl2fastq 2.17.1.14 (Illumina, Inc., 2017). Quality checks were performed on the raw data using FastQC v0.11.8 (Andrews, 2010) and MultiQC v1.7 (Ewels *et al*., 2016) using Python 3.6.8 (van Rossum and Drake, 2009). Due to differences in the number of reads for each library, raw FASTQ files were downsampled using seqtk 1.2 (Li, 2016) to the same depth of 1 million reads. Analysis of the fruit fly samples was performed with 1,000,000 reads for direct comparison with the plant samples.

Reads were trimmed using the FASTX-Toolkit 0.0.14 (Hannon, 2010) to remove the first eight bases as described in Townsley *et al*. (2015). All samples were trimmed to the same length for comparison purposes. Reads were filtered by quality using Trimmomatic 0.39 (Bolger *et al*., 2014) running on Java openjdk 1.8.0, because the algorithm supports processing of single and paired-end reads. All samples were filtered using the same quality threshold.

Samples were aligned to the respective reference genomes using Bowtie2 2.3.0 (Langmead and Salzberg, 2012) using default parameters. The TAIR10 reference for *Arabidopsis thaliana* was obtained from the Arabidopsis Information Resource database (Lamesch *et al*., 2011). The *Drosophila melanogaster* reference Berkeley Drosophila Genome Project assembly BDGP6.22 (dos Santos *et al*., 2015) was obtained from Ensembl release 96 (Cunningham *et al.*, 2019).

Coverage and summary statistics were computed using samtools 1.3.1 (Li *et al.*, 2009), bedtools 2.26.0 (Quinlan and Hall, 2010), and custom shell scripts. Data were processed and plotted using R 3.6.2 (R Core Team, 2019). The data.table 1.12.8 (Dowle and Srinivasan, 2019) and vroom 1.2.0 (Hester and Wickham, 2020) R packages were instrumental for processing large datasets efficiently. Library complexity was computed using preseq 3.0.2 (Daley and Smith, 2013). GC bias was computed using DeepTools v3.5.0 (Benjamini and Speed, 2012; Ramírez *et al.*, 2016)

Non-overlapping sliding (stepping) windows of 10,000 nucleotides and base frequencies at nucleotide resolution were computed using bedtools 2.26.0 (Quinlan and Hall, 2010). The coverage depth at the nucleotide resolution was computed using samtools 1.3.1 (Li *et al*., 2009) and sliding windows were computed in R 3.6.2 (R Core Team, 2019). Coverage depth was normalized by window length to account for truncated windows at the ends of chromosomes and scaffolds. The GC content was computed from base frequencies, and summary statistics for coverage depth were computed using GNU awk 4.1.4 (Aho *et al.*, 1988) and averaged in sliding windows in R 3.6.2 (R Core Team, 2019). Least-squares linear regression analysis was performed (y ~ β_1_x + β_0_), where y is the adjusted coverage depth and x is the GC content for each 10,000 bp sliding window for the genome. β_1_ is the coefficient for the magnitude of the linear relationship between the GC content and coverage depth.

## Results

By modifying the breath capture technology used in the BrAD-Seq protocol that was originally developed for RNA-seq, we succeeded in adapting the protocol for DNA sequencing (Fig. 1). To assess the utility of the novel protocol for genomic DNA sequencing, we compared its performance to the commercially available kits TruSeq (Illumina) and ThruPLEX (Rubicon Genomics) using plant *Arabidopsis thaliana* (*A. thaliana*) DNA and fruit fly *Drosophila melanogaster* (*D. melanogaster*) DNA. These kits have been shown to perform well with low DNA input (Sato *et al.*, 2019; Mauger *et al*., 2020) and are popular for WGS or ChIP-Seq analysis.

### BrAD-Seq generates libraries with high coverage breadth with less organelle genomes

Plant and fruit fly genomic DNA was processed using the BrAD-Seq protocol in duplicate and sequenced on the Illumina MiSeq and NovaSeq platforms, respectively. The raw reads were down-sampled to 1 million reads per sample to carry out comparisons (Fig. 2) with respect to the mapping rate of filtered reads (Fig. 2a and Fig. 2b), coverage fold-depth (mean read depth) (Fig. 2c and Fig. 2d), coverage breadth (proportion of the genome with three or more reads) (Fig. 2e and Fig. 2f), and uniquely mapped reads (Fig. 2g and 2h). Of note, due to poorer quality in Read 2s, both the mapping and unique mapping rates were lower with paired-end sequencing data (from flies) compared with single-end read sequencing data (plant, Fig. S1). Thus, fly data from Read 1 were analyzed further for direct comparison with the plant data. The proportion of mapped reads were similar between the raw data and the down-sampled sets (Tables S4 and S5), indicating the results are representative.

**Figure 2.**
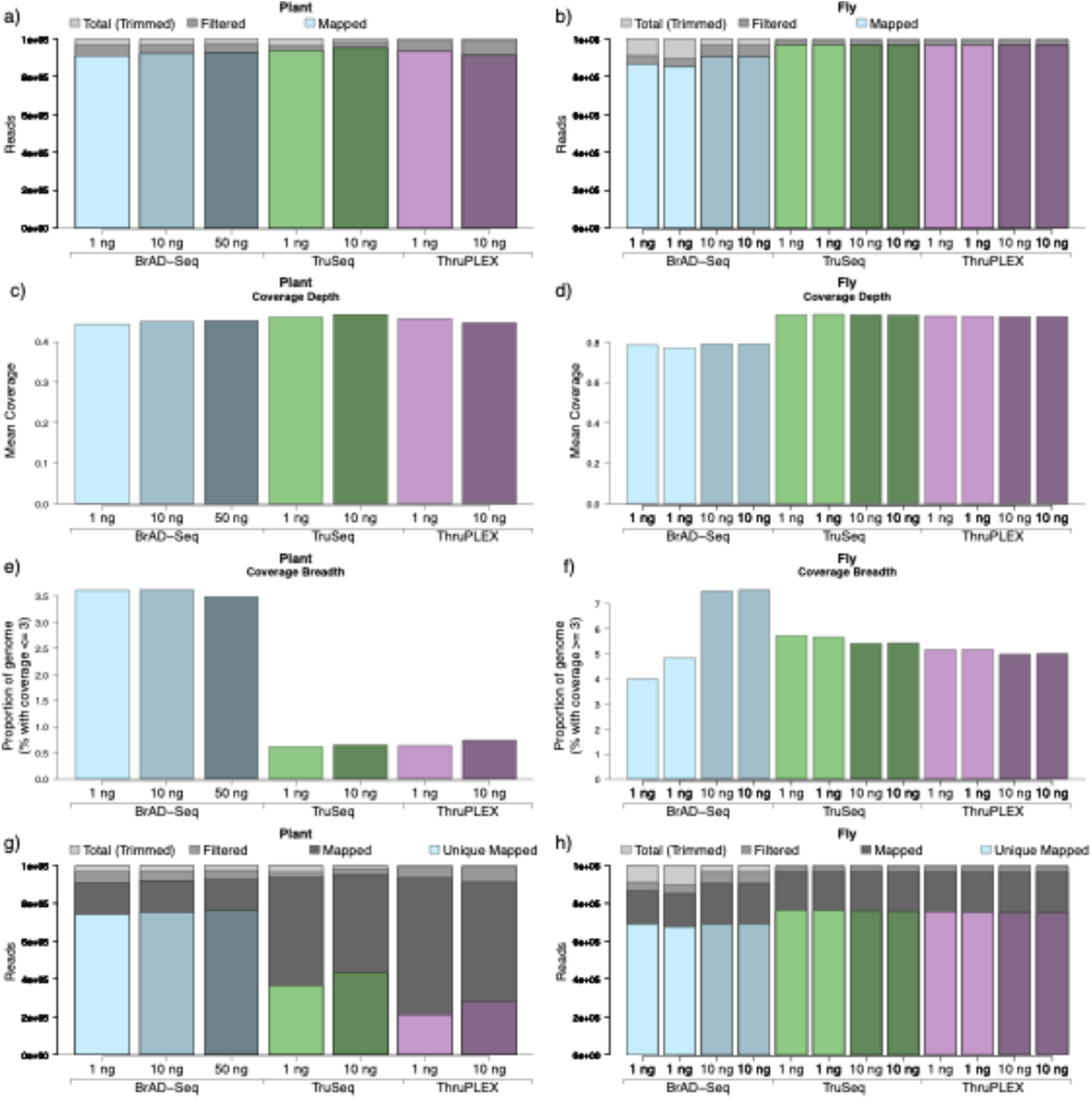
Data processing summary for different library preparation techniques. All libraries were downsampled to 1 million reads as single ends in read 1. The number of reads remaining after filtering and mapping to the reference genome is shown for a) plant data, and b) fly data after downsampling. Mapped reads are shown as a proportion of total trimmed reads with unmapped quality-filtered reads in dark grey, as described in the key. The mean coverage depth for plant and fly data closely reflects the number of mapped reads. The coverage breadth (the proportion of the genome mapped by ≥3 reads) for e) plant data, and f) fly data was often greater for many BrAD-Seq libraries. The unique mapping rate for g) plant data, and h) fly data shows that uniquely mapped reads were also higher in BrAD-Seq prepared using plant DNA and similar to other protocols using fly DNA. The fly samples were plotted in two replicates.

Our analysis shows BrAD-Seq libraries exhibit higher coverage breadth than the libraries prepared by other kits (Fig. 2e and Fig. 2f), indicating that BrAD-Seq libraries contain more sequences representing a wider range of genomic loci. This was more evident with the plant genomic DNA than with the fly genome (~3-fold higher for the plant and ~1.2-fold higher for the fly). In addition, BrAD-Seq libraries showed a higher unique mapping rate with the plant genomic DNA (Fig. 2g, h, and Table S4, S5). Therefore, BrAD-Seq appears to capture and cover more regions of the genome than other techniques, the beneficial effects of which appear to be more enhanced with the plant DNA. This was confirmed by determining proportions of mapped reads for each chromosome where we found that BrAD-Seq more accurately captured the expected ratio of mapped reads on *A. thaliana* chromosome 1 (87.1% of what is expected for chromosome length) compared with other techniques (34.9% in TruSeq; 31.7% in ThruPLEX, Fig. 3a). In addition, BrAD-Seq was less biased for multi-copy plastid genomes (Tables S6 and S7), exhibiting fewer mapped reads in plant chloroplasts compared with TruSeq (t = 6.97, p = 0.002) or ThruPLEX (t = 47.78, p = 4.38 × 10^−4^) and fly mitochondria compared with TruSeq (t = 73.17, p = 1.36 × 10^−12^) or ThruPLEX (t = 32.18, p = 9.48 × 10^−10^).

**Figure 3.**
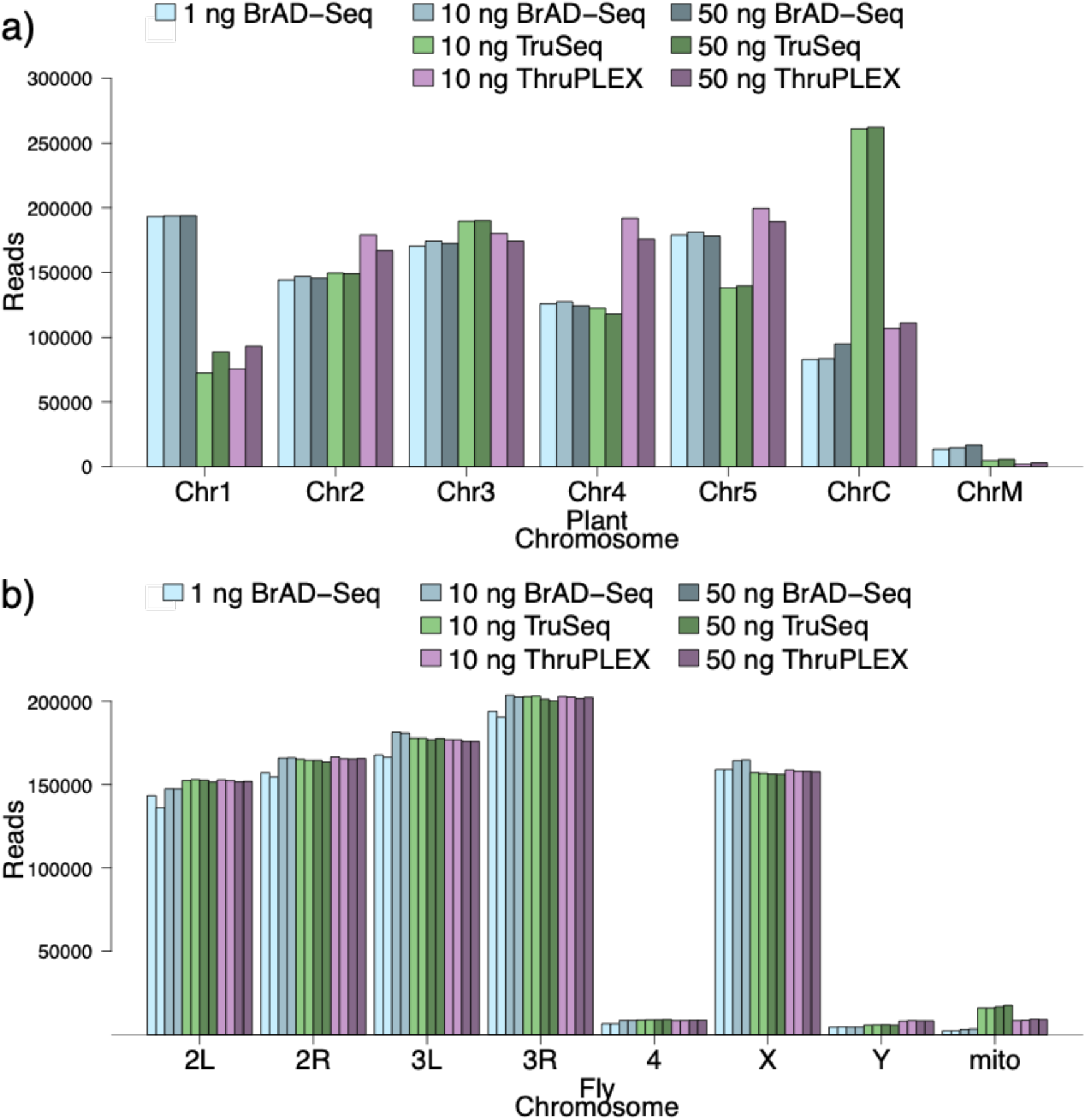
BrAD-Seq shows a relatively even distribution of reads per chromosome and for organelle genomes. All libraries were downsampled to 1 million reads, and the reads that mapped to each chromosome were counted. The breakdown of reads per chromosome for a) plant data, and b) fly data is shown. Unassembled scaffolds in the fly reference were excluded from the plot. In plants, BrAD-Seq resulted in more reads for the longest chromosome (1), and TruSeq libraries showed over-representation of chloroplast reads. Both TruSeq and ThruPLEX libraries contained more mitochondrial reads (Fig. S2 for details).

### BrAD-Seq shows tolerance for variability in GC content

We next examined library complexity by resampling raw data and computing the number of unique molecules obtained (using preseq). Whereas the input amount of plant genomic DNA had a relatively small impact on library complexity, clear differences were observed between the protocols (Fig. 4a). The BrAD-Seq samples had a higher library complexity, which may be explained by the higher coverage breadth (Fig. 2e) and unique mapping rates (Fig. 2g). However, this was not replicated with the fly genomic DNA, which unexpectedly showed low library complexity (Fig. 4b).

**Figure 4.**
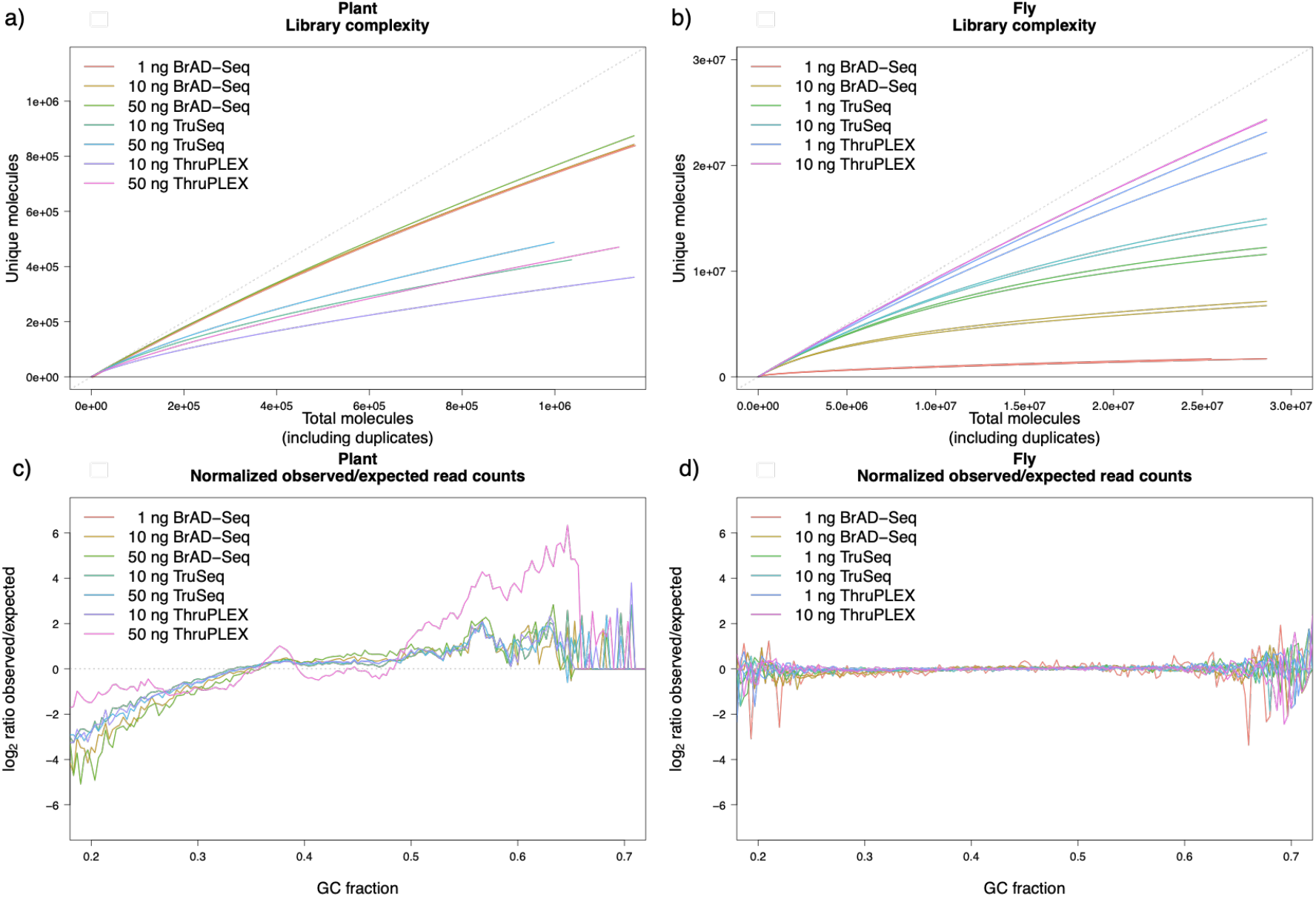
Library complexity and coverage bias across the genome. Library complexity is shown by plotting unique molecules by resampling the raw unmapped reads as computed by preseq 3.0.2 for a) plant samples, and b) fly samples. Library complexity was higher in BrAD-Seq for plant samples at all input amounts but was lower in fly samples in the various replicates, especially for 1 ng of BrAD-Seq libraries. GC bias, as computed by DeepTools 3.5.0, is shown for 300 bp fragments in c) plant, and d) fly samples. Illumina and Rubicon libraries showed a greater bias towards high GC regions in the plant genome. A straight line at 0 indicates no GC bias compared to the expected distribution of sampling of the reference genome. Some variations at extreme values are expected because of sampling a smaller number of fragments.

In respect to the GC content, the plant libraries had a higher GC content than the genome average (36%) with TruSeq libraries showing higher GC content and variance than others (Table S8). The fly libraries had a similar mean (42%) and standard deviation to the genome (Table S9). GC bias was computed from aligned reads using DeepTools v3.5.0 for a fragment size of 300 bp (Benjamini and Speed, 2012; Ramírez *et al.*, 2016). Compared to the statistically expected distribution, BrAD-Seq libraries exhibited less bias towards high GC regions of the plant genome (Fig. 4c), but all techniques showed less bias in the fly samples (Fig. 4d).

Regions with high GC content can be biased towards higher coverage depths due to lower annealing temperatures. Therefore, a linear relationship is expected between coverage and GC content for GC-biased techniques (Chen *et al.*, 2013). Such bias was more pronounced in TruSeq and ThruPLEX libraries than in BrAD-Seq, and BrAD-Seq libraries did not show a linear trend or elliptical distribution indicative of GC bias (Fig. 5). This result is supported by the lower effect size of the GC content with BrAD-Seq compared to others measured by the slope (β_1_) of the least-squares linear regression analysis (Table 2). Taken together, these results show that the BrAD-Seq protocol has less bias for GC content compared to the currently available low-input Illumina TruSeq and Rubicon ThruPLEX library preparation kits.

**Table 2.** GC coverage bias across sliding windows of 10,000 bp in *A. thaliana* and *D. melanogaster*.

| Library | Slope ($\beta_1$) in plants | Slope ($\beta_1$) in flies |
| --- | --- | --- |
| 10 ng BrAD-Seq | 3.6874 | 62.114 |
| 10 ng TruSeq | 9.2336 | 179.720 |
| 10 ng ThruPLEX | 6.0395 | 148.933 |

\* $\beta_1$ is the coefficient for linear regression for $y \sim \beta_1 x + \beta_0$ , where y is the coverage depth and x is the GC content for each 10,000 bp sliding window of the genome.

**Figure 5.**
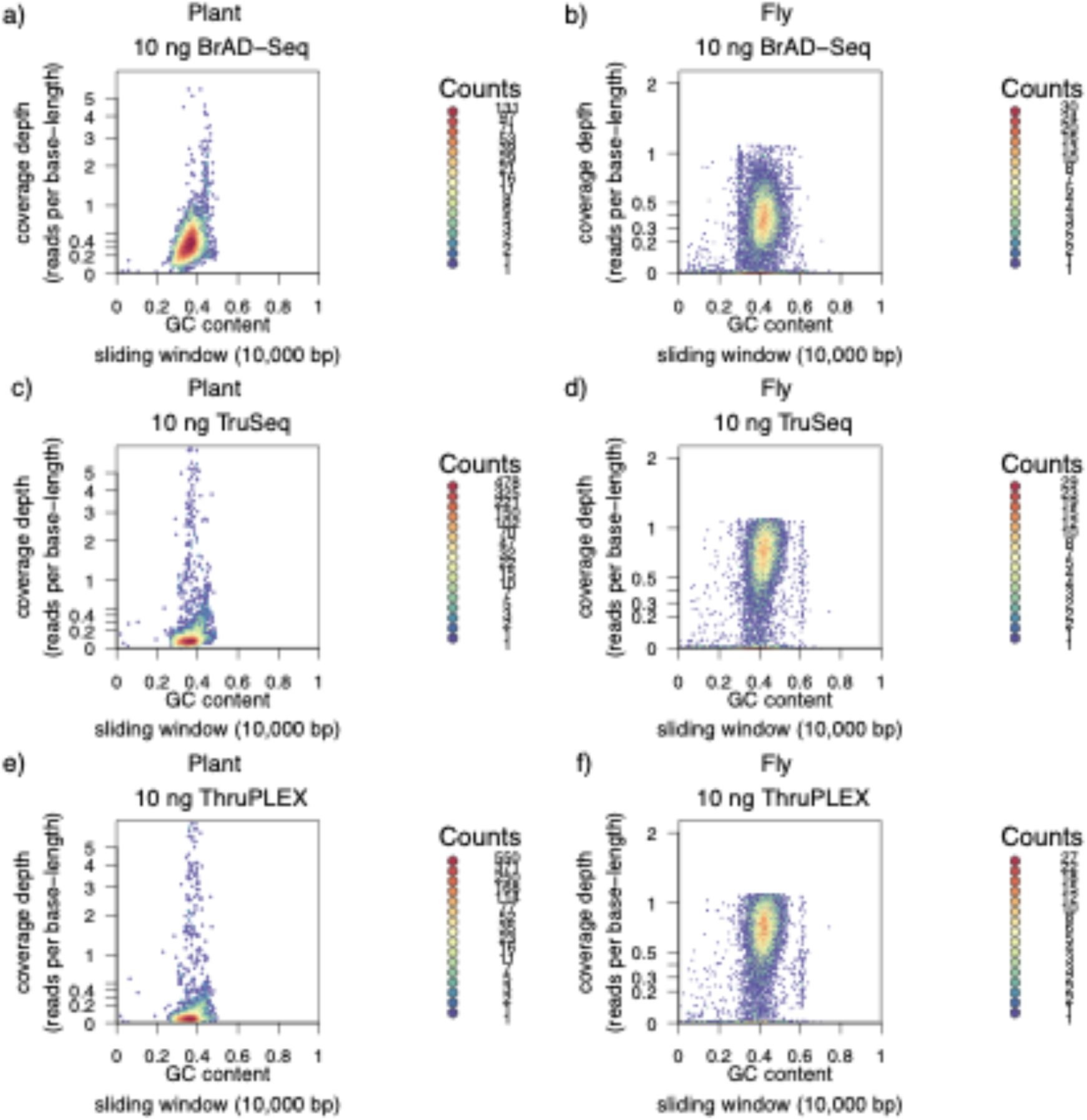
Coverage depth in sliding windows is associated with GC content in biased libraries. Coverage depth varies with GC content at fine physical resolution, and the relationship differs between libraries. Reads were binned into 10,000 bp stepping windows in the genome and plotted on a log-scale against the GC base frequency for the same regions in the genome. Overlapping points were binned into hexagons and colored by density. Examples of a-b) BrAD-Seq, c-d) TruSeq, and e-f) ThruPLEX libraries are shown for 10 ng DNA input in plant (a, c, and e) and fly (b, d, and f) samples. TruSeq and ThruPLEX libraries had similar distributions and showed bias towards fragments with high GC content, particularly in plant samples. Summary statistics for GC content are presented in Tables S8 and S9. The linear regression results (Table 21) show that there was a greater linear effect of GC content on the read depth in the TruSeq or ThruPLEX libraries. In the low GC plant genome, Illumina and Rubicon libraries exhibited bias for GC-rich regions at a finer window resolution.

## Discussion

Here, we present a fast, simple, and low-cost novel library preparation protocol for genomic DNA that can be used with low input amounts. The BrAD-Seq protocol is compatible with the Illumina sequencing platform and can be used for a wide range of species and applications. The BrAD-Seq protocol provides higher quality libraries, which are less biased towards multi-copy plastids and GC-rich regions of the genome than commercially available kits specific for low DNA inputs. The novel method appears to be especially advantageous for sequencing analysis for plants (e.g., *A. thaliana*), based on observations of higher unique mapping rates, less enrichment for plastid genes, and low bias for fragments with high GC content. It should also be readily applied to epigenomic analyses such as ChIP-Seq, chromosome conformation capture, and DNA footprinting, which tend to yield low amounts of DNA. Because sequence quality was higher for single-end sequencing than paired-end reads in the BrAD-Seq libraries due to the lower quality of read 2, single-end sequencing is recommended although paired-end sequencing can still be performed. It is of note that BrAD-Seq libraries appear to have lower read quality and mapping rates; however, sequencing at a sufficient depth can overcome this drawback and is arguably offset by reduced coverage bias, fast turn-around time, and reduced consumables costs.

To summarize, we present the BrAD-Seq protocol which has a higher coverage breadth, library complexity, and low GC bias as demonstrated in plant samples. Therefore, BrAD-Seq addresses the need for a low-cost, low-input technique with low GC bias in plants and other species with low GC content in the genome. By reducing costs (Lakens *et al.*, 2021), the novel method improves the possibility of carrying out genomics studies with a larger number of samples in a diverse range of species.

## Data Availability

All sequencing data were deposited in the NCBI sequence read archive (SRA) BioProject <u>PRJNA735014</u> and <u>PRJNA735016</u> for *Arabidopsis thaliana* and *Drosophila melanogaster*, respectively.

## Acknowledgments

We thank Dr. Shunsuke Ishii (RIKEN CPR) for providing us with *D. melanogaster* genomic DNA (DGRC #5905). Sequencing was performed at the RIKEN IMS internal facilities and was outsourced to Takara Bio Japan K.K. We thank Dr. Shungo Kobori for his advice on the data analysis. This work was supported by a JSPS KAKENHI Grant-in-Aid for Scientific Research on Innovative Areas “Principles of pluripotent stem cells underlying plant vitality” (JP17H06470) to AM, the Cabinet Office, Government of Japan, Cross-ministerial Moonshot Agriculture, Forestry and Fisheries Research and Development Program, “Technologies for Smart Bio-industry and Agriculture” (funding agency: Bio-oriented Technology Research Advancement Institution) to YI.

## Author’s contributions

TH, KH, and YI developed and optimized the novel experimental technique to prepare BrAD-Seq libraries. HO-Y and SK prepared the libraries using the commercial kits. STK, AM, and YI designed the experiments and prepared the manuscript. STK performed all data processing and bioinformatic analyses. AM and YI oversaw the project.

